# ALKALs are *in vivo* ligands for ALK family Receptor Tyrosine Kinases in the neural crest and derived cells

**DOI:** 10.1101/210096

**Authors:** Andrey Fadeev, Patricia Mendoza Garcia, Uwe Irion, Jikui Guan, Kathrin Pfeifer, Stephanie Wiessner, Fabrizio Serluca, Ajeet Pratap Singh, Christiane Nüsslein-Volhard, Ruth H. Palmer

## Abstract

Mutations in Anaplastic Lymphoma Kinase (ALK) are implicated in somatic and familial neuroblastoma, a paediatric tumour of neural crest-derived tissues. Recently, biochemical analyses have identified secreted small ALKAL proteins (FAM150, AUG) as potential ligands for human ALK and the related Leukocyte Tyrosine Kinase (LTK). In the zebrafish *Danio rerio*, DrLtk, which is similar to human ALK in sequence and domain structure, controls the development of iridophores, neural crest-derived pigment cells. Hence, the zebrafish system allows studying Alk/Ltk and Alkals involvement in neural crest regulation *in vivo*. Using zebrafish pigment pattern formation, *Drosophila* eye patterning, and cell culture-based assays, we show that zebrafish Alkals potently activate zebrafish Ltk and human ALK driving downstream signalling events. Overexpression of the three Dr Alkals cause ectopic iridophore development whereas loss of function alleles lead to spatially distinct patterns of iridophore loss in zebrafish larvae and adults. *alkal* loss of function triple mutants completely lack iridophores and are larval lethal as is the case for *ltk* null mutants. Our results provide the first *in vivo* evidence of (i) activation of ALK/LTK family receptors by ALKALs and (ii) an involvement of these ligand-receptor complexes in neural crest development.

## Introduction

Receptor tyrosine kinases (RTK) constitute a large family of surface receptors involved in a wide range of cellular processes, both in normal development and carcinogenesis. RTK inhibitors show a strong promise as a targeted patient-specific therapy (Murphy et al., 2017). Despite intense interest in RTKs, two of them remain obscure in terms of their biochemistry and native function - Anaplastic Lymphoma Kinase (ALK) and the related Leukocyte Tyrosine Kinase (LTK) which constitute a subgroup of receptor tyrosine kinases (ALK RTKs) involved in human cancers (Hallberg and Palmer, 2013).

Human ALK (HsALK) has a unique extracellular domain (ECD) composition among receptor tyrosine kinases containing two MAM domains (after meprin, A-5 protein and receptor protein-tyrosine phosphatase µ), an LDLa domain (Low-density lipoprotein) and a glycine-rich domain (GR). In comparison, the extracellular domain of human LTK (HsLTK) is smaller and lacks the LDLa and both MAM domains (Figure 1A) (Hallberg and Palmer, 2013). Although ALK ligands have been identified in invertebrates, no evidence has been presented describing ligand activation of vertebrate ALKs *in vivo*.

**Figure 1.**
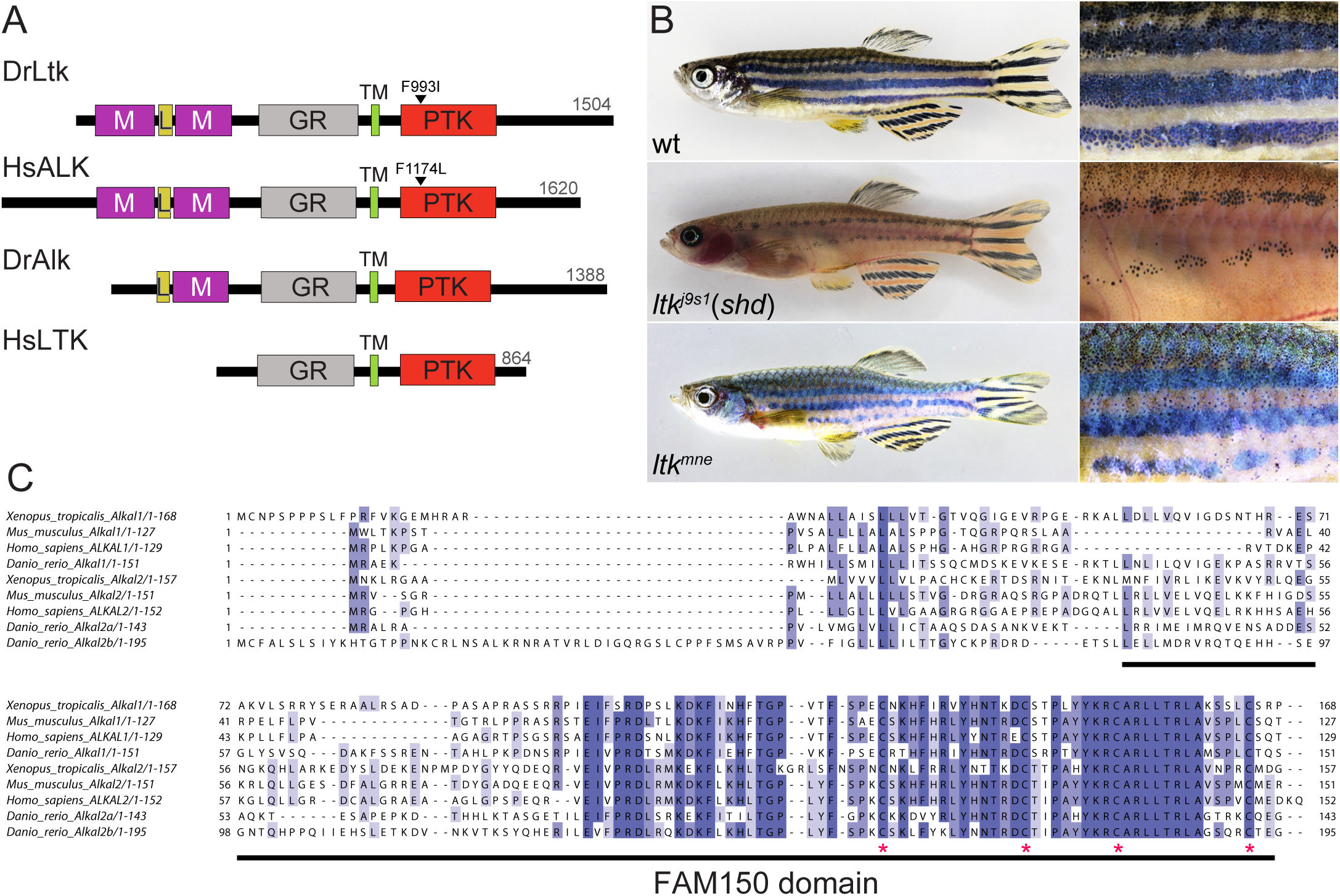
Ectopic expression of ALKALs and overactivation of DrLtk leads to supernumerary iridophores in *D. rerio.* A) Domain structure of the human and zebrafish ALK/LTK RTK family. M: MAM, L: LDLa, GR: glycine rich, TM: transmembrane domain, PTK: kinase domain. The activating F993I (DrLtk) and F1174L (HsALK) mutations within the kinase domain are indicated. (B) *ltk* loss-of-function *shady* mutants (*ltk^j9s1^*) lack iridophores, while gain-of-function *moonstone* (*ltk*^*mne*^) exhibit increased numbers of iridophores. (C) Alignment of ALKAL proteins from different species. Underlined - the FAM150 domain, red asterisks - conserved Cys. Note high conservation of the C-terminal half of the FAM150 domain.

While knowledge of endogenous ALK and LTK function in humans is limited, ALK, as a result of protein fusion, overexpression or activating mutations, is involved in the development of various cancer types (Hallberg and Palmer, 2013). Mice lacking ALK exhibit defects in neurogenesis and testosterone production but are viable, as are *ALK;LTK* double mutants (Bilsland et al., 2008; Weiss et al., 2012; Witek et al., 2015). The *D. melanogaster* and *C. elegans* genomes each contain a single ALK RTK – *DAlk* and *sdc-2* respectively (Lorén et al., 2001; Reiner et al., 2008). Zebrafish (*D. rerio*) has two members of the ALK RTK family – Ltk (DrLtk) and Alk (DrAlk) (Lopes et al., 2008; Yao et al., 2013). DrLtk possesses two MAM domains, and thus resembles HsALK while DrAlk has a smaller ECD that lacks one MAM domain (Figure 1A, Figure 1-figure supplement 1) (Lopes et al., 2008). Mammalian *LTK* is expressed in pre-B and B lymphocytes and brain (Ben-Neriah and Bauskin, 1988; Bernards and de la Monte, 1990) whereas zebrafish *alk* is expressed in the developing central nervous system (Yao et al., 2013). In contrast, both zebrafish *ltk* and mammalian *ALK* are expressed in neural crest (Iwahara et al., 1997; Lopes et al., 2008; Vernersson et al., 2006). Expression pattern and domain structure suggests that HsALK is more similar to DrLtk than DrAlk (Figure 1A, Figure 1-figure supplement 1).

In zebrafish, Ltk has a firmly established role in the development of iridophores in pigment pattern formation: mutants in *shady* (*shd*), the gene encoding DrLtk, lack iridophores and display patterning defects (Fadeev et al., 2016; Frohnhöfer et al., 2013; Lopes et al., 2008) (Figure 1B). Zebrafish develops two pigment patterns throughout its life. The first is a larval pattern that derives directly from migrating neural crest cells during early embryogenesis. This simple pattern is comprised of relatively evenly distributed yellow xanthophores and three stripes of black melanophores, accompanied by silvery iridophores lining the dorsal and ventral stripes. Dense sheets of iridophores cover the iris of the eye and the swim bladder (Figure 2C) (Kelsh et al., 1996; Singh and Nüsslein-Volhard, 2015). The adult zebrafish pattern of longitudinal light and dark stripes develops during metamorphosis (3-6 weeks post fertilisation) (Figure 1B). Adult iridophores arise from stem cells located in the dorsal root ganglia; these stem cells are multipotent and produce all three pigment cell types, as well as peripheral neurons and glia (Singh et al., 2014, 2016). Iridophores appear in the skin along the horizontal myoseptum where they proliferate and spread dorso-ventrally to successively form the light stripes by patterned aggregation (Fadeev et al., 2015; Frohnhöfer et al., 2013; Patterson and Parichy, 2013; Singh and Nüsslein-Volhard, 2015; Singh et al., 2014). Numerous iridophores cover the iris of the eyes, the operculum of the gills and the exposed margins of the scales. Proliferation and spreading of iridophores is controlled by homotypic competition (Walderich et al., 2016). Interestingly, while iridophores are indispensable for striped pattern formation in the trunk, they are not required for stripe formation in the fins (Fadeev et al., 2015; Frohnhöfer et al., 2013; Singh et al., 2014, 2016).

**Figure 2.**
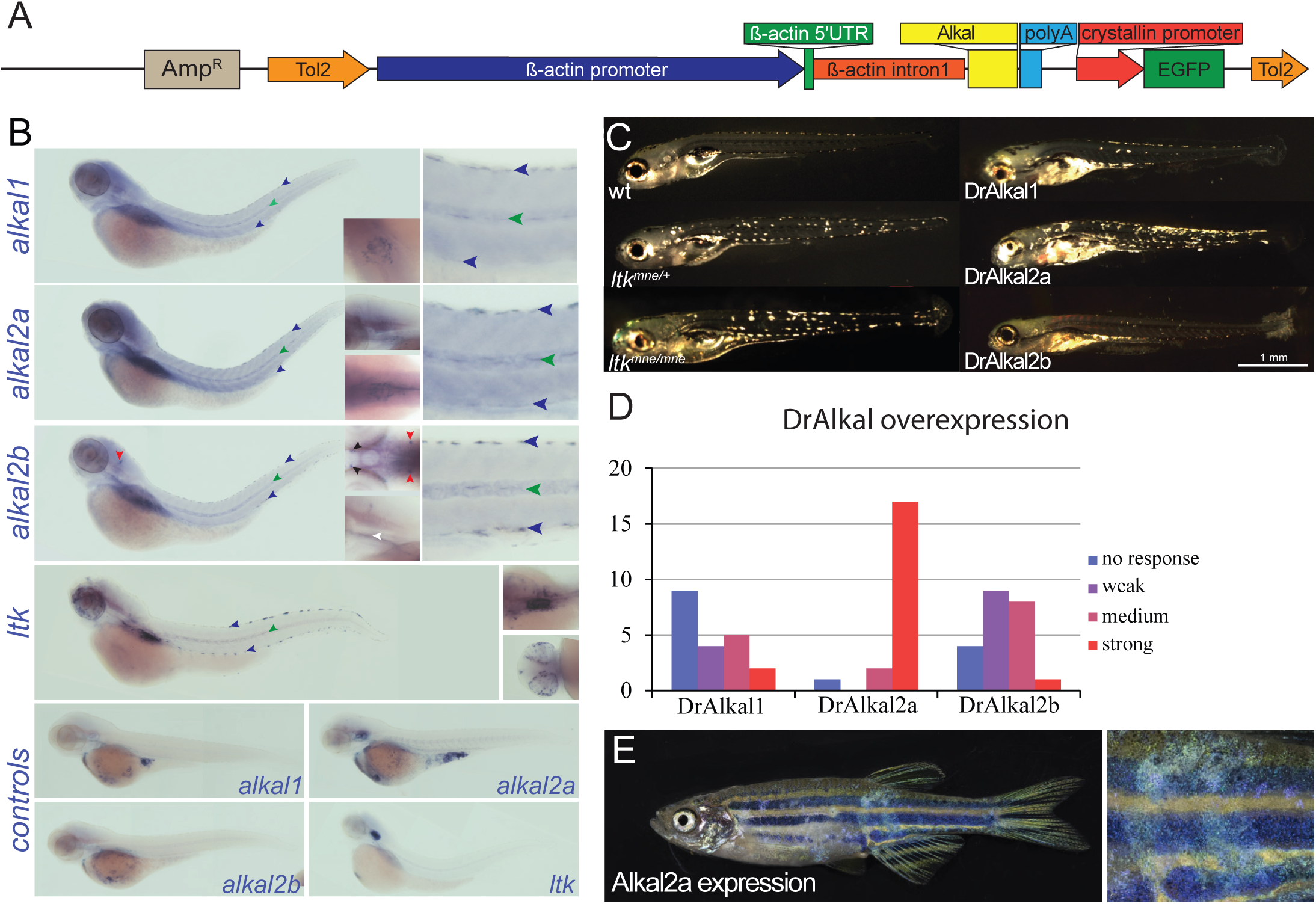
Expression of DrAlkals is sufficient for ectopic production of iridophores. (A) Schematic of *ALKAL*-expression constructs. Amp^R^ – ampicillin resistance, Tol2 – medaka Tol2 transposase recognition sequence. (B) *alkal* expression in 72 hpf zebrafish larvae. Weak *alkal1* signal can be observed in the head, swim bladder, notochord (green arrowheads) and iridophores in dorsal and ventral stripes (blue arrowheads). *alkal2a* mRNA is observed in the head, swim bladder, notochord (green arrowheads) and iridophore stripes (blue arrowheads). *alkal2b* mRNA is detected in swim bladder, and a row of cells at the ventral aspect of the head (white arrowhead), bilateral clusters in the head (red arrowheads), behind the eyes (black arrowheads), notochord (green arrowheads) and iridophores stripes (blue arrowheads). *ltk* expression is visible in swim bladder, eyes, notochord and iridophores. Negative controls: corresponding sense probes. (C) Supernumerary iridophores were observed in 5 dpf larvae upon ectopic expression of indicated DrAlkals in F0 injected fish. *ltk*^*mne*^ larvae are shown as control. (D) Distribution of phenotypes of (1C). (E) Mosaic overexpression of *alkal2a* produces patches of supernumerary iridophores in adults.

In *shd* mutants iridophores are missing; complete loss-of-function in *ltk* leads to larval lethality, whereas mutants carrying weaker alleles are adult viable (Figure 1B) (Frohnhöfer et al., 2013; Lister et al., 1999; Lopes et al., 2008; Parichy et al., 2000). *ltk* is expressed in the early neural crest and gradually becomes restricted to iridophores (Lopes et al., 2008). Chimeras, obtained by blastomere transplantations, revealed that *ltk* is autonomously required in iridophores (Carney et al., 2008; Frohnhöfer et al., 2013). Taken together, these findings lead to the conclusion that *ltk* is required for the establishment and proliferation of iridophores and their progenitors from multipotent neural crest cells (Lopes et al., 2008).

Recently, a gain of function zebrafish mutant *moonstone* (*ltk*^*mne*^), harbouring a hyperactive point mutation in Ltk (F993I), analogous to the human ALK-F1174 hotspot observed in neuroblastoma patients, was described (Fadeev et al., 2016). Point mutations are the most common cause of ALK activation in both familial and sporadic neuroblastoma, and the corresponding mutation in HsLTK, F568L, has been shown to cause ligand-independent activity (Carén et al., 2008; Chand et al., 2013; Chen et al., 2008; George et al., 2008; Janoueix-Lerosey et al., 2008; Mossé et al., 2008; Roll and Reuther, 2012). Fish carrying the *ltk*^*mne*^ mutation display ectopic iridophores in the trunk as larvae, and an increased number of iridophores on scales and fins as adults, giving them a strong blue-green sheen (Figure 1B). Moreover, treatment with the ALK inhibitor TAE684 during metamorphosis stages decreases the number of iridophores, suggesting that Ltk-activity is continually required for iridophore development and maintenance. When presented with a wildtype environment in chimeric animals, *ltk*^*mne*^ iridophores can massively overgrow in the surrounding skin, suggesting that Ltk is involved in control of iridophore proliferation by homotypic interactions (Fadeev et al., 2016; Walderich et al., 2016).

The identity of the ligand(s) for vertebrate ALK and LTK has been elusive for many years, and although ligands for ALK have been identified in invertebrates – Jeb in *Drosophila* and HEN-1 in *C. elegans* – homologous ligands in vertebrates remain unidentified (Englund et al., 2003; Ishihara et al., 2002; Lee et al., 2003; Reiner et al., 2008; Weiss et al., 2001). Recently human secreted small proteins ALKAL1 and 2 (for “ALK And LTK ligand”, HsALKALs), which were previously reported as family-with-sequence-similarity-150 (FAM150) or Augmentor (AUG), have been shown to activate human ALK and LTK in cell culture and when co-expressed in *Drosophila* eyes (Guan et al., 2015; Reshetnyak et al., 2015; Zhang et al., 2014). Characterised by the highly conserved FAM150 domain at their C-terminus and four conserved cysteines (Figure 1C), HsALKALs bear no obvious similarities to Jeb and HEN-1. In contrast to the two HsALKALs, the zebrafish genome harbours three ALKAL homologues (DrAlkals), encoded by *alkal1*, *alkal2a* and *alkal2b*. Hereafter we will refer to HsALKALs and DrAlkals together as ALKALs. No physiological function has been ascribed to ALKALs so far in any model organism, nor has *in vivo* genetic evidence supporting activation of ALK RTK signalling by ALKALs been presented. The structural similarities together with the shared neural crest pattern of tissue expression between zebrafish Ltk and human ALK offer an opportunity to define the role of the ALKAL ligands in activation of the ALK family of RTKs *in vivo*.

In this study, we present evidence for an *in vivo* interaction between ALKALs and DrLtk, a member of the vertebrate ALK RTK family. Local overexpression of the DrAlkals results in excess iridophore production in larvae, and an iridophore overgrowth in the adult striped pattern. We show in mammalian cell cultures and in *Drosophila* eyes that DrLtk and HsALK are activated by zebrafish and human ALKALs (ALKALs). HsALK activation by the zebrafish Alkal2a in human neuroblastoma cells demonstrates evolutionary conservation of this interaction. Similar to *ltk* loss-of-function mutants, knockout of the zebrafish *alkal* genes reduces iridophore numbers, indicating that these ligands act in a spatially specific, partially redundant manner. The triple mutant, in which all three Alkals are knocked out using the CRISPR/Cas9 system, is larval lethal and displays a complete loss of iridophores as does a *ltk* null allele. This demonstrates that all three Alkals are necessary to fully activate DrLtk in target tissues. The activation of DrLtk and HsALK by human and zebrafish ALKALs provides the first genetic evidence for the interaction of ALKALs and ALK/LTK RTKs as an evolutionarily conserved functional signaling unit in the neural crest and neural crest-derived tumours.

## Results

### DrAlkals and DrLtk have similar expression domains

The zebrafish genome harbours three FAM150 domain containing ALKAL homologues (DrAlkals), encoded by *alkal1*, *alkal2a* and *alkal2b* (Figure 1C). We employed *in situ* probes against *alkal1*, *alkal2a* and *alkal2b* to characterise their larval expression patterns (Figure 2B). *alkal1* signal is weak but detectable in the iridophore stripes, the eye primordium and the swim bladder. *alkal2b* expression is wider but strong in the swim bladder and single cells of unknown identity in the head. The strongest signal was observed for *alkal2a*, expressed in the notochord and iridophore stripes of the trunk, as well as in the eye and swim bladder. In agreement with previous data (Lopes et al., 2008), we observed *ltk* mRNA in the head and iridophores of the trunk and eyes, as well as in the swim bladder (Figure 2B). Thus, *alkals* appear to be expressed in a pattern overlapping with that of *ltk*.

### Overexpression of ALKALs results in ectopic iridophores in zebrafish

To investigate the effect of ALKALs on iridophore development we used a genome integration cassette driving Alkal expression under the ubiquitous β*-actin* promoter to drive mosaic overexpression (Figure 2A). Clonal expression of all three zebrafish Alkals resulted in the appearance of ectopic iridophores in 5 days post fertilization (dpf) larvae, similar to *ltk*^*mne*^ mutants (Figure 2C). The phenotypes were classified into four classes by the number of ectopic iridophore clusters (or single cells): 1. no response, 2. weak (<10), 3. medium (10-30) and 4. strong (> 30) phenotypes (Figure 2 – figure supplement 1A). While Alkal1- and 2b- overexpressing larvae exhibited mostly no, weak or medium phenotypes, expression of Alkal2a produced predominantly strong responses (Figure 2D). As adults, Alkal2a overexpressing fish exhibited patches of supernumerary iridophores on scales, fins and eyes – a phenotype similar to that of *moonstone* mutants (Figure 2E). Strikingly, frequently patches of dense clusters of tumour-like disorganised iridophore overgrowth appeared in the skin resembling the iridophore clusters in chimeras in which *ltk*^*mne*^ iridophores develop in juxtaposition with *ltk*^*+*^ iridophores. Ectopic iridophores could also be induced by human and mouse ALKALs (Figure 2 – figure supplement 1B). Taken together these results suggest that overexpression of ALKALs leads to the activation of endogenous ALK RTKs *in vivo*.

### Human and zebrafish ALKALs activate zebrafish Ltk in the *Drosophila* eye

We next analysed the activation of ALK/Ltk receptors by Alkal proteins by coexpressing them in *Drosophila* eyes, taking advantage of the clear read-out of this system. In brief, ligands and receptors were ectopically expressed in the fly eye using the *GAL4-UAS* system (Brand and Perrimon, 1993) employing *pGMR-GAL4* to drive expression in the developing fly eye. Activation of the ALK/LTK receptors in this assay results in a rough eye phenotype (Guan et al., 2015; Martinsson et al., 2011). Both human ALKALs activated HsALK and lead to a rough eye phenotype, in agreement with previous data (Guan et al., 2015)(Figure 3A). As with the HsALK receptor, both HsALKALs robustly activated DrLtk killing flies raised at 25°C. Weaker transgene expression, achieved by rearing flies at 18°C, resulted in severe rough eye phenotypes (Figure 3A). No phenotype was observed when wildtype Ltk was expressed alone (Figure 3A). In contrast, expression of the constitutively activate Ltk^mne^ led to a rough eye phenotype (Figure 3 - figure supplement 1A). Ectopic expression of either of the zebrafish Alkals together with DrLtk (but not alone) resulted in rough eyed flies, thus all three DrAlkals are able to activate DrLtk (Figure 3B). These data suggest the ability of ALKAL proteins to activate ALK/LTK receptors is conserved across vertebrates.

**Figure 3.**
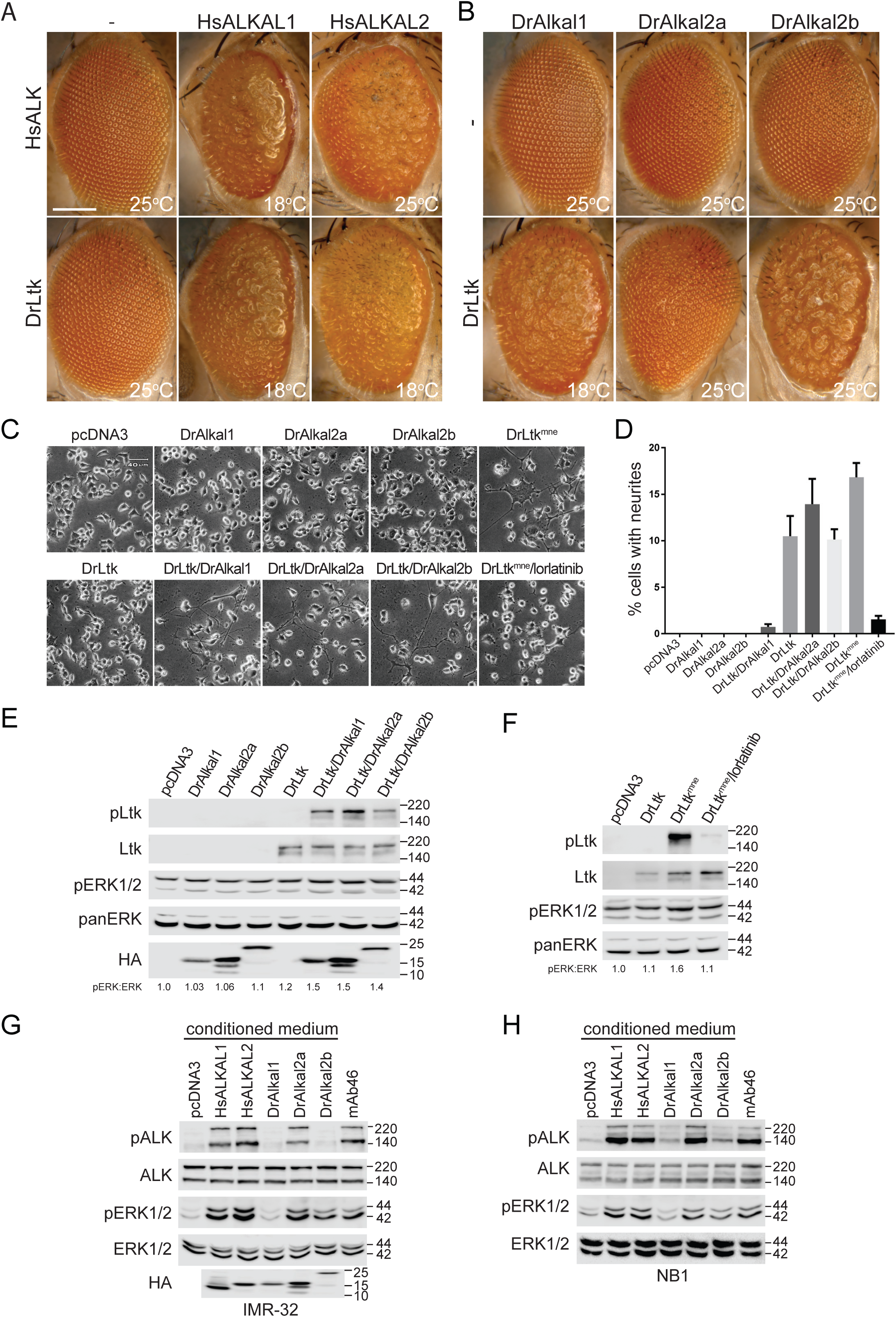
Human and zebrafish ALKALs activate DrLtk and HsALK. (A) Ectopic expression of either HsALK or DrLtk in combination with HsALKALs, but not alone, leads to a disorganization of the ommatidial pattern in *Drosophila* eye. (B) Expression of DrAlkals and DrLtk together, but not alone, results in rough eye phenotypes. Temperatures at which flies were grown are indicated. (C) Neurite outgrowth in PC12 cells expressing either *pcDNA3* vector control, DrAlkals or DrLtk alone, combinations of DrLtk and DrAlkals, or DrLtk^mne^ in the presence or absence of the ALK tyrosine kinase inhibitor lorlatinib as indicated. (D) Quantification of neurite outgrowth in PC12 cells as indicated in (C). Error bars: SD. (E) Immunoblot of whole cell lysates from PC12 cells expressing DrLtk and DrAlkals alone or in combination as indicated. (F) Immunoblot of whole cell lysates from PC12 cells expressing wildtype DrLtk and DrLtk^mne^ in the presence or absence of lorlatinib as indicated. (G,H) Activation of endogenous HsALK by ALKAL-containing medium in neuroblastoma cell lines: IMR-32 (G) and NB1 (H). Antibodies against ALK, ERK and actin were used as loading controls. pALK-Y1278 - phosphorylated HsALK or DrLtk. α-myc - total DrLtk protein. HA antibodies - ALKAL proteins. pERK1/2 indicates activation of downstream signalling. Agonist mAb46 ALK was employed as positive control.

### ALKALs activate zebrafish Ltk and human ALK

To investigate activation of DrLtk by the ALKAL proteins in more detail and in a more controlled environment, we employed a rat PC12 cell neurite outgrowth assay. Expression of zebrafish DrAlkals or DrLtk alone did not lead to neurite differentiation, unlike Ltk^mne^, which promoted neurite extension and ERK activation downstream of Ltk in the absence of ALKALs (Figure 3C-E). This effect was abrogated by the addition of the ALK inhibitor lorlatinib, shown to inhibit the effect of human ALK-F1174 activating mutations, analogous to *ltk*^*mne*^ (Guan et al., 2016; Infarinato et al., 2016)(Figure 3C, D, F). Co-expression of zebrafish Alkals with DrLtk also led to robust induction of neurite outgrowth, phosphorylation of the Ltk receptor and ERK activation, in agreement with our findings in the *Drosophila* eye assay (Figure 3 C-E). Furthermore, DrLtk was activated by both human ALKALs (Figure 3 – figure supplement 1B,C) whereas DrAlkal2a (but not 1 or 2b) led to robust neurite outgrowth when expressed together with HsALK (Figure 3 – figure supplement 1D). Thus, this independent assay also supports ALK RTKs activation by ALKALs in a manner dependent on the tyrosine kinase activity of the receptor.

### Endogenous HsALK signalling in neuroblastoma cells is activated by zebrafish Alkal2a

To address whether endogenous levels of human ALK respond to DrAlkal proteins we employed DrAlkal-conditioned medium to stimulate IMR-32 neuroblastoma cells that express wildtype ALK, using phosphorylation of ALK and ERK as a read-out. As shown previously, ALK was robustly activated by both HsALKALs (Figure 3G, Figure 3 – figure supplement 1D) (Guan et al., 2015). In agreement with our results in PC12 cells, endogenous HsALK was visibly activated by Alkal2a, but not Alkal1 or Alkal2b, suggesting that these proteins may be less effective in activating human ALK as well as the DrLtk receptor in larvae (Figure 3G). Similar results were obtained in the NB1 neuroblastoma cell line, where *ALK* lacks exon 2-3 (Figure 3H). In summary, using multiple approaches, we demonstrate that ALKALs are able to activate zebrafish Ltk and human ALK.

### Regulation of iridophores by DrAlkal requires Ltk

To test whether DrLtk is critical for DrAlkal-dependent induction of iridophores in zebrafish we investigated the effect of DrAlkal2a overexpression in fish lacking *ltk*. To do this we generated fish carrying knockout frameshift mutations in *ltk* using the CRISPR/Cas 9 system to introduce 2 bp and 22 bp insertions in the first exon. *ltk* knockout (*ltk*^*ko*^) larvae did not develop any iridophores (Figure 4A). Similarly, larvae treated with the ALK inhibitor lorlatinib did not develop iridophores, consistent with the results of neurite outgrowth assays (Figure 4B). When overexpressed in *ltk*^*ko*^ mutants, DrAlkal2a was no longer able to produce ectopic iridophores, whereas expression of DrAlkal2a in heterozygous or wildtype siblings led to increased number of iridophores as observed earlier (Figure 4A). This indicates that DrAlkal2a acts upstream of DrLtk to generate iridophores. The *ltk*^*ko*^ mutant fish did not survive past 6 dpf; and lethality may be in part due to failure of the swim bladder to fill with air. Interestingly, the strong adult viable *shady* allele *ltk*^*j9s1*^ was still able to produce ectopic iridophores, suggesting the production of a normally inactive, but still ligand-responsive Ltk^j9s1^ protein in these *ltk* (*shd*) mutants (Figure 4A). We also observed that the hyperactive *ltk*^*mne*^ allele can be further activated by ectopic expression of DrAlkal2a, which shows dense iridophore patches in the striped pattern, overgrowing the dark melanophore stripe, a phenotype never seen in *ltk*^*mne*^ individuals (Figure 4C). These results support the role of DrAlkals as effectors of iridophore development acting specifically through the DrLtk receptor.

**Figure 4.**
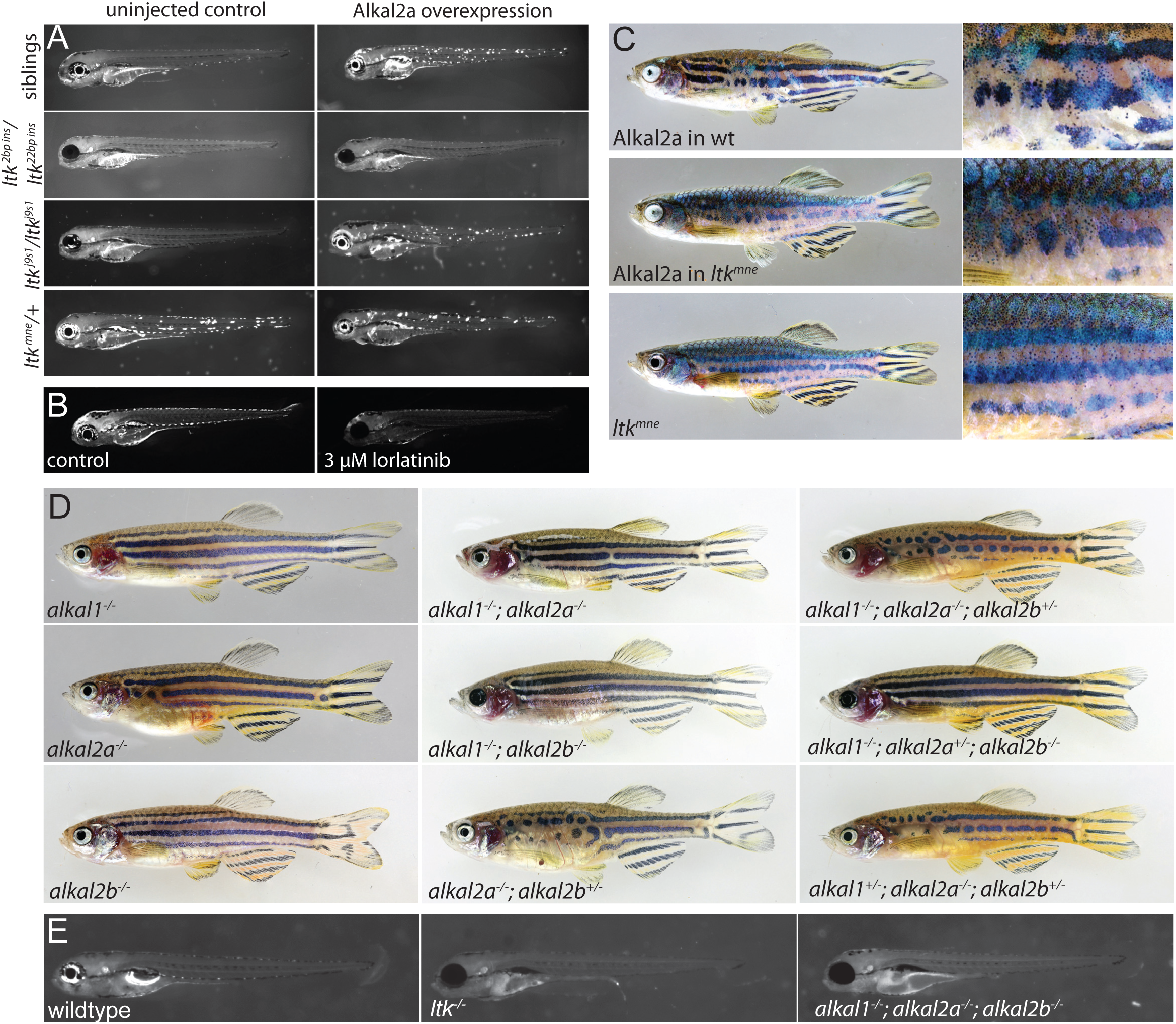
DrAlkals affect iridophore development via DrLtk. (A) Ectopic expression of DrAlkal2a does not rescue loss of iridophores in 4 dpf transheterozygous *ltk* knockout zebrafish larvae (2 bp insertion/22 bp insertion, n=9/9), whereas in heterozygous or wildtype siblings and *ltk*^*j9s1*^ larvae it leads to overproduction of iridophores (n=29/29 and 34/35). Overexpression of DrAlkal2a in *ltk*^*mne*^ mutant slightly enhances the phenotype. (B) Treatment with lorlatinib, results in diminished iridophore numbers in larvae. (C) Mosaic overexpression of DrAlkal2a produces patches of supernumerary iridophores both in wildtype and *ltk*^*mne*^adults. Overexpression of DrAlkal2a in *ltk*^*mne*^ mutants enhances production of supernumerary iridophores compared to wildtype. (D) Zebrafish *alkal* mutants display defects in iridophore development. *alkal1*^*ko*^ mutants have reduced iridophores in eyes and operculum, and removal of both *alkal1*and *alkal2b* results in a complete loss of eye iridophores. *alkal2a*^*ko*^ mutants show the strongest phenotype, with reduced numbers of iridophores in the trunk, especially anteriorly. This phenotype displays increased penetrance in *alkal2a;alkal2b* double mutants. (E) Triple mutant for all *alkal* genes is embryonic lethal and displays total loss of iridophores, similar to *ltk* transheterozygous knockout.

### Loss of DrAlkals leads to spatially specific loss of iridophores

To better understand the function of endogenous DrAlkals and further strengthen evidence of ALK RTK activation by ALKALs *in vivo*, we knocked out DrAlkals utilizing CRISPR/Cas9 technology. Many F0 mosaic loss-of-function fish (especially those mutant for *alkal2a*) exhibited patches devoid of iridophores during metamorphosis, which in most cases disappeared by adulthood, presumably due to proliferation of residual iridophores successively filling the space devoid of iridophores (Walderich et al., 2016). Since presence of iridophores during metamorphosis is crucial for correct stripe formation in the trunk (but not fins), we investigated the iridophores and stripe pattern of adult mutants, completely lacking functional Alkals. *alkal1* mutants exhibit strong reduction of iridophore number exclusively on the operculum with no detectable phenotype in the trunk (Figure 4D). *alkal2a* mutant fish show reduced dense iridophores in the light stripes of the trunk, particularly in the anterior region, resulting in irregular interruptions of melanophore stripes (Figure 4D). *alkal2b* mutant fish, in contrast, do not exhibit any detectable trunk phenotype in adults. The *alkal1* mutant phenotype is further enhanced by mutation in either *alkal2a* or *alkal2b*. For example, mutants in both *alkal1* and *alkal2b*, lack iridophores on the operculum and in the eye. The *alkal2a;alkal2b* double mutant is larval lethal with mutants failing to inflate the swim bladder. The *alkal1;alkal2a;alkal2b* triple mutant is also larval lethal and displays a larval phenotype indistinguishable from *ltk* knockout alleles (Figure 4E). The trunk phenotype of *alkal2a* mutants is enhanced by additionally removing one copy of *alkal2b,* which on its own does not show a detectable phenotype. Interestingly, of the six copies representing the three *alkal* genes, one copy of *alkal2a* or *alkal2b* alone is sufficient to produce viable fish with a reduced number of iridophores and some patterning defects, indicating that the requirement of Alkal protein for activation of DrLtk is very low (Figure 4D). In summary, we observe overlapping and partially redundant spatial functions of the three Alkals in zebrafish.

## Discussion

This work provides the first *in vivo* evidence that the ALKAL (FAM150/Augmentor) class of ligands bind and activate the ALK receptor tyrosine kinase. We show that overexpression of DrAlkals produce responses in zebrafish similar to overactivation of DrLtk (*moonstone*) – namely supernumerary and ectopic iridophores. Likewise, loss of function of these proteins results in severe reductions of iridophore numbers, both in larvae and in adults. Human and zebrafish ALKALs activate zebrafish Ltk signaling in both, *Drosophila* eye and neurite outgrowth assays, in a manner dependent on the tyrosine kinase activity of the receptor. In addition, zebrafish Alkal2a and human ALKALs strongly activate exogenous human ALK in a neurite outgrowth assay and the endogenously expressed ALK receptor in neuroblastoma cells.

We demonstrate the involvement of a new ligand-receptor interaction in the control of neural crest-derived iridophores. Importantly this Ltk-Alkal interaction controls both larval iridophores directly differentiating from neural crest, and metamorphic iridophores, which come from multipotent stem cells and produce the adult pigmentation pattern (Singh and Nüsslein-Volhard, 2015; Singh et al., 2014, 2016). The three ligands display slightly different spatial requirements, but with the exception of the iridophores on the operculum, which absolutely require Alkal1, other regions of the body are devoid of iridophores only when the function of more than one Alkal is eliminated by mutation. The complete loss of function of all three Alkals results in the total absence of iridophores and in larval lethality, a phenotype indistinguishable from that of Ltk mutants. This indicates that the three Alkals cooperate to fully activate the receptor and ensure the full repertoire of iridophore development.

Our expression analysis shows that in the zebrafish embryo, *alkals* are expressed in the regions where iridophores develop, and where also Drltk is expressed. This raises the possibility that in this system ligand and receptor are expressed in the same cells, presumably in the iridophores themselves. Other ligand-receptor pairs have already been suggested to play important roles in zebrafish pigmentation. *Endothelin receptor b1* (*rose*) and *endothelin 3b* are suggested to function in iridophore cell shape transitions (Fadeev et al., 2016; Krauss et al., 2014), *kita* (*sparse*) and *kit ligand a* (*kitla* or *sparse-like*) are necessary for establishing embryonic and most adult melanophores. Xanthophores are controlled by homotypic competition and depend on Csf signaling, however *kitla* and *csf1*-ligands are not expressed in corresponding pigment cells, i.e., melanophores and xanthophores respectively (Dooley et al., 2013; Patterson and Parichy, 2013; Walderich et al., 2016). The differences in expression pattern of different ligand-receptor pairs may have implications on the dynamics of color pattern formation in zebrafish and related *Danio* species. Our analysis suggests that Alkal/DrLtk mediated signaling triggers iridophore proliferation in zebrafish, where this signaling pathway might link homotypic competition (known to regulate iridophore proliferation) to cell cycle machinery. Co-expression of ligand and receptor in the same cells may lead to autocrine/paracrine activation of Ltk, resulting in the high rate of proliferation of iridophores observed during metamorphosis in zebrafish. The function of Ltk signaling therefore might be to stimulate cell proliferation, differentiation and survival rather than providing guidance cues for directed migration, e.g. as a morphogen gradient. However, it is likely that there are additional constraints on iridophore proliferation during development – while metamorphic iridophores tend to spread where possible and cover the whole skin, larval iridophores do not do so. Ectopic expression of Alkal ligands in zebrafish causes increased numbers of iridophores in larvae as well as adult fish, suggesting that the effect of Alkal/DrLtk signaling on iridophore proliferation is instructive. In summary, color pattern formation in zebrafish offers an exciting opportunity to investigate this question valuable to our understanding of the fundamental role of ALK in cell proliferation and cancer.

It appears that zebrafish Ltk displays more similarities to the human ALK RTK than to LTK. These similarities are not only in the ECD domain structure, but importantly also include expression in the neural crest. This raises an intriguing question of orthology relationship between human and zebrafish genes. While the original designation of Ltk in zebrafish was based upon similarities in the kinase domains, based upon analysis of the entire proteins DrLtk is more similar to HsALK than HsLTK (Figure 1 - figure supplement 1). It is feasible that HsALK and DrLtk are orthologues, and that DrAlk possibly arose by duplication from DrLtk and that a more correct designation would be DrLtk-> DrAlk1, and DrAlk->DrAlk2. Future characterization of the zebrafish ALK family members should help resolve this at the functional level. In the meantime, this work provides not only genetic and biochemical evidence for the ALKAL/ALK family RTK interaction, but also demonstrates conservation of this interaction in vertebrates, since human ligands are able to activate zebrafish receptors and vice versa. Furthermore, we show the importance of this interaction for the regulation of neural crest derived cells. This becomes especially significant when the role of HsALK-F1174 gain-of-function mutations in human neuroblastoma is compared to the effect of the gain-of-function mutation in the exactly the same position in DrLtk^mne^ (F993), which leads to ectopic differentiation of neural crest-derived iridophores. Our findings, and their significance, are further corroborated by complementary work recently published by (Mo et al., 2017). We hope that the present study will facilitate further investigations in the role of ALKALs in neuroblastoma. Outstanding questions remain, such as the individual contribution of the zebrafish Alkals in terms of spatial and temporal regulation of Ltk signaling, that should be addressed in future studies. In summary, this work confirms a conserved role for the ALKAL ligands in activation of ALK family RTKs in the developing neural crest.

## Materials and Methods

### Zebrafish maintenance and image acquisition

The following fish lines were bred and maintained as described (Nüsslein-Volhard and Dahm, 2002): TÜ wildtype, *shady*^*j9s1*^ (Lopes et al., 2008), *ltk*^*mne*^ (Fadeev et al., 2016). All experiments with zebrafish were performed in accordance with the guidelines of the Max-Planck-Society and approved by the Regierungspräsidium Tu□bingen, Baden-Wu□rttemberg, Germany (Aktenzeichen: 35/9185.46). Anesthesia was performed as described previously (Singh et al., 2014). Canon 5D Mk II and Leica M205FA were used to obtain images. Adobe Photoshop and Illustrator CS6 and 4 were used for image processing. For adult fish photos multiple RAW camera images were taken in different focal planes and auto-align and auto-blend functions of Photoshop were used. Blemishes on the background were removed using the brush tool, without affecting the image of the fish.

### Multiple alignments and trees

Multiple protein alignment was obtained using MUSCLE (Edgar, 2004) and visualized using Jalview (Waterhouse et al., 2009). Neighbour joining was used to construct the trees (Saitou and Nei, 1987). The following GenBank sequences were used for analysis: LTK - ACA79941.1 (*Danio rerio*), P29376 (*H. sapiens*); ALK - XP_691964.2 (*D. rerio*), NP_004295.2 (*H. sapiens*); ALKAL1 - NP_001182661.1 (*M. musculus*), NP_997296.1 (*H. sapiens*), XP_012820744.1 (X. tropicalis), XP_005166986.1 (D. rerio); ALKAL2: NP_001153215.1 (M. musculus), NP_001002919.2 (H. sapiens), XP_004914540.1 (X. tropicalis), XP_015140421.1 (G. gallus), XP_002665250.2 (Alkal2b, D. rerio), XR_659754.3 (Alkal2a, D. rerio, translated from 1771-2202 bp).

### Injections

Microinjections into zebrafish eggs were performed as described (Nüsslein-Volhard and Dahm, 2002). Zebrafish Injection Pipettes OD 20µm (BioMedical Instruments, Germany) and Pneumatic PicoPump SYS-PV820 (World Precision Instruments, Sarasota, Florida) were used with the following parameters: pressure 30 psi, period 1.0s.

### Overexpression and knockout

The following mixes were used to inject yolk of one cell stage embryos: for knockout - 350ng/µl Cas9 protein, 15ng/µl sgRNA and 0.05% Phenol Red (P3532, Sigma-Aldrich, St. Louis, Missouri); for ectopic expression of DrAlkal: 12.5 ng/µl Tol2 mRNA (Kawakami and Shima, 1999), 25ng/µl of a plasmid and 0.05% Phenol Red. ALKAL-overexpression plasmids were provided by FS and SW. Cas9 protein was expressed in *E.coli* (BL21 DE3 pLysS) with an aminoterminal Twin-Strep-Tag and 6xHis tag. The protein was purified according to the manufacturer’s instructions (IBA lifesciences, ‘Expression and purification of proteins using double tag [Strep-tag/6xHistidine-tag]’ vPR36-0001) and dialysed into PBS +300 mM NaCl+100 mM KCl. sgRNAs targeting zebrafish *alkal* coding sequences were prepared as described before (Irion et al., 2014), using the following primers:

*alkal1*_sgRNA_for TAGGTCTGCTGTTGTCCGGCTT

*alkal1*_sgRNA_rev AAACAAGCCGGACAACAGCAGA

*alkal2a*_sgRNA_for TAGGACACGCACCATCTCAAAA

*alkal2a*_sgRNA_rev AAACTTTTGAGATGGTGCGTGT

*alkal2b*_sgRNA_for TAGGAGCCCTATGAAGACAGGT

*alkal2b*_sgRNA_rev AAACACCTGTCTTCATAGGGCT.

### Lorlatinib treatment

Lorlatinib (PF 06463922, Tocris, Bio-Techne, Minneapolis, MN, USA) was dissolved in dimethylsulphoxide (DMSO; Sigma-Aldrich Louis, MI, USA) to a final concentration of 0.1 mM. Standard E2 embryonic medium (Nüsslein-Volhard and Dahm, 2002) was supplemented with 50 μM gentamycin (NH09, Carl Roth GmbH + Co. KG, Karlsruhe, Germany) and 0.03% DMSO with 3 μM lorlatinib or without for controls. Thirty embryos were manually dechorionated at 9 hpf and treated in 50 ml of the medium in a 30 mm 1% agarose coated Petri dish with daily changes of medium for fresh one.

### Genotyping of zebrafish

DNA from embryos or adult caudal fin biopsies was prepared as described in (Meeker et al., 2007). The regions surrounding CRISPR-targets were amplified with RedTaq DNA Polymerase (D4309, Sigma-Aldrich) and sequenced using Big Dye Terminator v3.1 kit (4337455, Thermo Fisher Scientific, Waltham, Massachusetts) with the following primers:

*alkal1*_for AGCAAGGAGGTAAAGGAGTC

*alkal1*_rev CACTCTTTTGATCTACAGAGGG

*alkal2a*_for TGGGGCTCGTATTGTTAATC

*alkal2a*_rev GCATAACAGAGTACACCCCA

*alkal2b*_for TGCTTTGCGTTATCGTTATCA

*alkal2b*_rev AGACTAGCAGGGAGTCAGCG.

### Whole mount in situ hybridization

Partial cDNA sequences (Ensembl v10) of zebrafish *alkals* and *ltk* cloned into pcDNA3 were used as templates to generate sense and antisense probes using SP6- and T7-Megascript Kits (Ambion/Lifescience). Detailed sequences are shown in Supplementary File 1. The whole mount *in situ* hybridization was carried out after Pfeifer et al 2012. Embryos were treated with 10 µg/ml Proteinase K for 10 min.

### Expression of Ltk and Alkal in *Drosophila* eye

*Drosophila* were maintained on a potato-mash diet under standard husbandry procedures. Crosses were performed at 25°C unless otherwise stated. *UAS- ltk*, *UAS- ltk^mne^, UAS-alkal1a, UAS-alkal2a and UAS-alkal2b* were synthesised (GenScript, Piscataway, New Jersey) and transgenic flies were obtained by injection (BestGene Inc., Chino Hills, California). *white*^*1118*^ (*w^1118^*) flies were employed as control. *GMR-Gal4* fly stock (Ref. #1104) was from the Bloomington *Drosophila* Stock Center at Indiana University (BDSC; NIH P40OD018537). *UAS-ALK*, *UAS-ALK^F1174L^*, *UAS-ALKAL1* and *UAS-ALKAL2* were generated previously (Guan et al., 2015). Fly eye samples were analysed under Zeiss Axio Imager.Z2 and AxioZoom.V16 microscopes. Images were acquired with an Axiocam 503 colour camera.

### Generation of DrLtk and DrAlkal expression constructs

Zebrafish *ltk*, *ltk*^*mne*^, *alkal1*, *alkal2a* and *alkal2b* constructs were synthesised in pcDNA3 vector (Genscript, Piscatay, NJ, USA). *ltk* constructs were tagged C-terminally with myc, while Alkal constructs were tagged C-terminally with HA. Human ALK and ALK-F1174L have been described previously (Martinsson et al., 2011).

### Neurite outgrowth assay

PC12 cells (2×10^6^) were co-transfected with 0.5 μg of pEGFP-N1 vector (Clontech, Mountain View, CA, USA) together with one or more of the following constructs: 0.5 μg of pcDNA3- *HsALK,* 1.0 μg of pcDNA3 empty vector,1.0 μg of pcDNA3 vector containing zebrafish *alkal1*, *alkal2a*, *alkal2b*, *ltk*, *ltk*^*mne*^, or human *HsALKALs*. Cells were transfected using an Amaxa Nucleofector electroporator (Amaxa GmbH, Cologne, Germany) with Ingenio electroporation solution (Mirus Bio LCC, Madison, WI, USA) and transferred to RPMI1640 medium (HyClone, GE Healthcare Bio-Sciences Austria GmbH, Pasching, Austria) supplemented with 7% horse serum (Biochrom AG, Berlin, Germany) and 3% FBS (Sigma-Aldrich Chemie GmbH, Steinheim, Germany). Approximately 5% of cells were seeded into 12-well plates for neurite outgrowth assays. The remaining cells were seeded in 6-well plates for immunoblotting assays. The ALK inhibitor lorlatinib, was used at a final concentration of 30nM to inhibit the activity of Ltk^mne^ (Guan et al., 2016). Two days after transfection, the percentage of GFP-positive and neurite-carrying cells versus GFP-positive cells was analysed under a Zeiss Axiovert 40 CFL microscope. Cells with neurites longer than twice the length of the cell body were considered neurite carrying. Experiments were performed in triplicate and each sample within an experiment was assayed in duplicate.

### Stimulation of endogenous HsALK with human and zebrafish ALKALs

ALKAL-containing conditioned medium was generated as described previously (Guan et al., 2015). In brief, HEK293 cells in complete DMEM medium (HyClone, GE Healthcare Bio-Sciences Austria GmbH, Pasching, Austria) were seeded into 10-cm dishes and transfected with 6 μg of pcDNA3 empty vector, *ALKAL1*, *ALKAL2*, *alkal1*, *alkal2a* or *alkal2b*- codingplasmid using lipofectamine 3000 (Invitrogen, ThermoScientific, Waltham, MA, USA). Medium was changed to neuroblastoma cell medium (RPMI1640, 10% FBS) 6 hours after transfection, and conditioned medium was collected after another 30-40 hours. Human neuroblastoma IMR-32 or NB1cells seeded in 12-well plates were stimulated with conditioned medium or the agonist monoclonal antibody mAb46 for 30 minutes prior to lysis (Moog-Lutz et al., 2005).

### Cell lysis and immunoblotting

Electroporated PC12 cells seeded in 6-well plates as described above were cultured in RPMI1640 medium supplemented with 7% horse serum and 3% FBS for 24 hours and then starved for 36 hours. Both PC12 cells and human neuroblastoma cells were lysed directly in 1x sodium dodecyl sulfate (SDS) sample buffer. Precleared lysates were run on SDS-PAGE. DrLtk or HsALK phosphorylation was analysed with pALK-Y1278 antibodies (Cell Signaling Technology, Leiden, The Netherlands) and activation of downstream components was detected with α-pERK1/2 antibodies (Cell Signaling Technology, Leiden, The Netherlands). Pan-ERK (BD Biosciences, San Jose, CA, USA) and/or β-actin (13E5) (Cell Signaling Technology, Leiden, The Netherlands) were employed as loading control. Total Ltk was detected with anti-Myc antibody (ab9132, Abcam, Cambridge, UK). Total ALK was detected with ALK (D5F3) antibody (Cell Signaling Technology, Leiden, The Netherlands). ALKALs were detected with anti-HA.11 antibodies (Clone 16B12, Biolegend, San Diego, CA, USA).

## Author contribution

AF and UI: zebrafish experiments, except *in situ* performed by KP. JG, PM and RHP: cell culture and *Drosophila* eye experiments. SW and FS: initial observations of ALKALs overexpression in zebrafish. AF, RP and APS: study design. AF, RP and CNV: writing the manuscript. All authors were involved in the final version of the manuscript.

## Figure legends

**Figure 1 - Figure Supplement 1.**
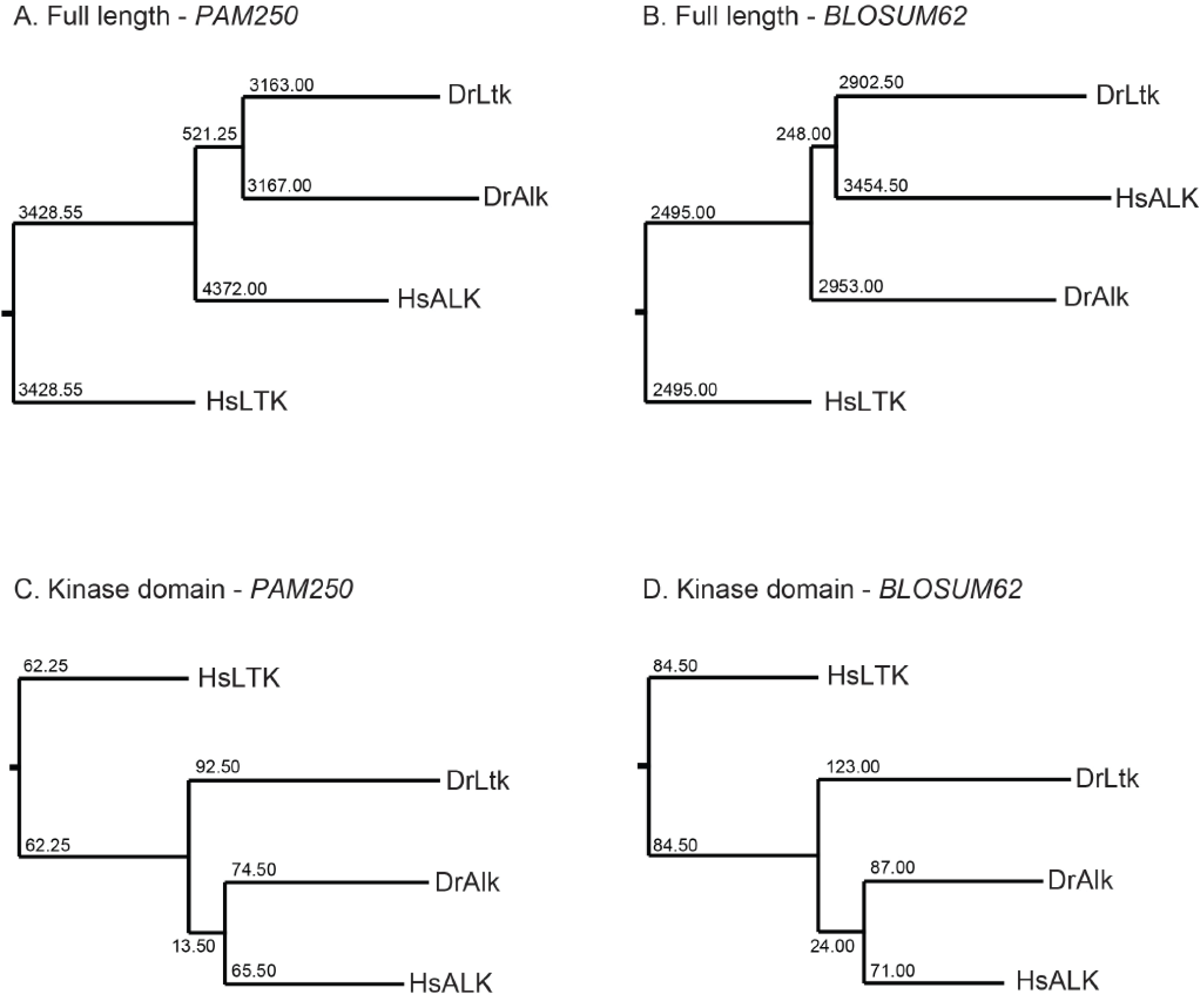
DrLtk is more similar to HsALK than to HsLTK. Analysis of protein sequences similarities using PAM250 (A,C) or BLOSUM62 (B,D) matrices on full length protein sequences (A,B) or sequences of kinase domains (C,D). Numbers indicate distances.

**Figure 2 - Figure Supplement 1.**
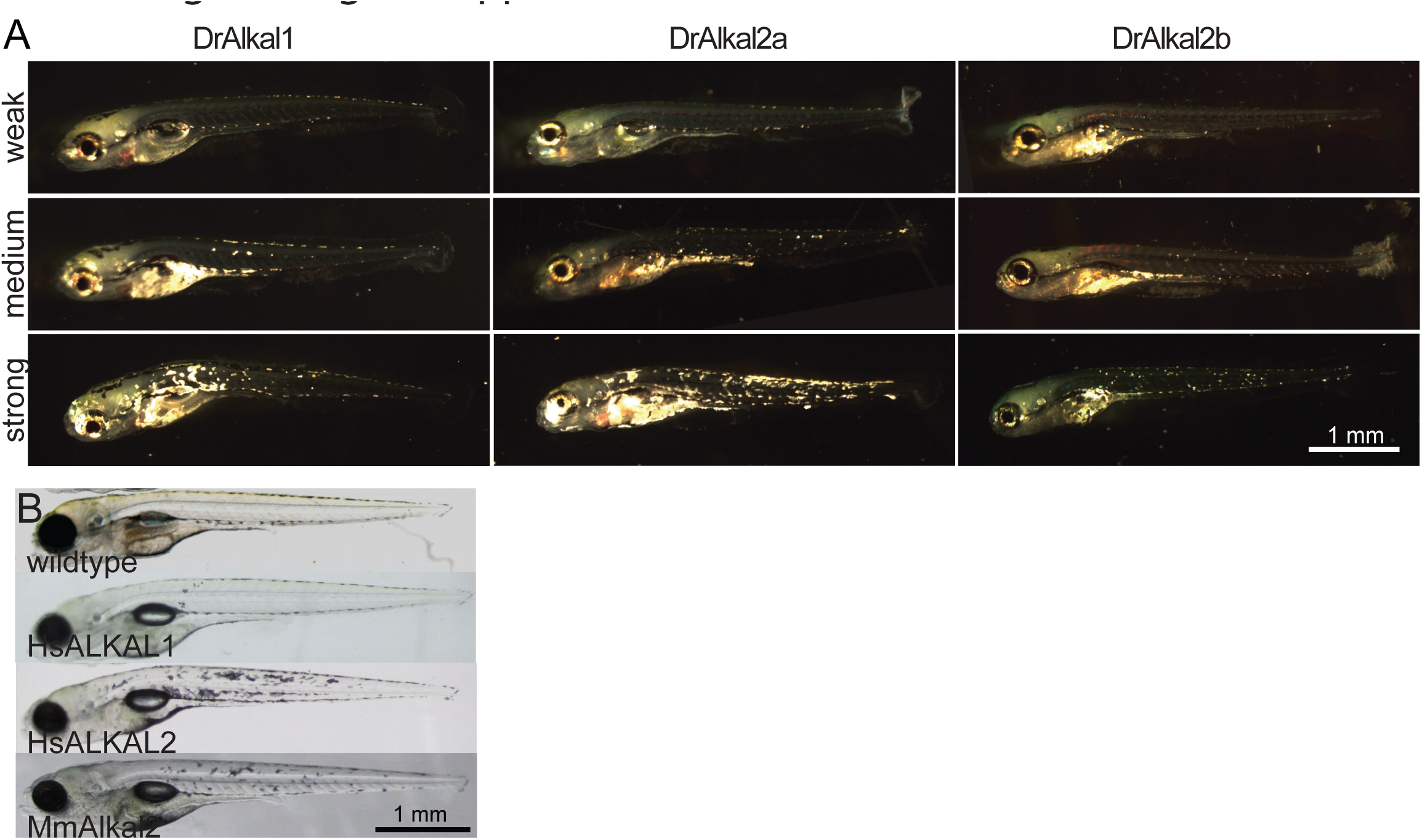
Ectopic iridophores in 5 dpf zebrafish larvae. Ectopic iridophores observed in 5 dpf larvae injected with constructs driving expression of zebrafish Alkals (A) or human and mouse ALKALs (B). Shown are examples of weak, medium and strong phenotypes, as used in Figure 1.

**Figure 3 - Figure Supplement 1.**
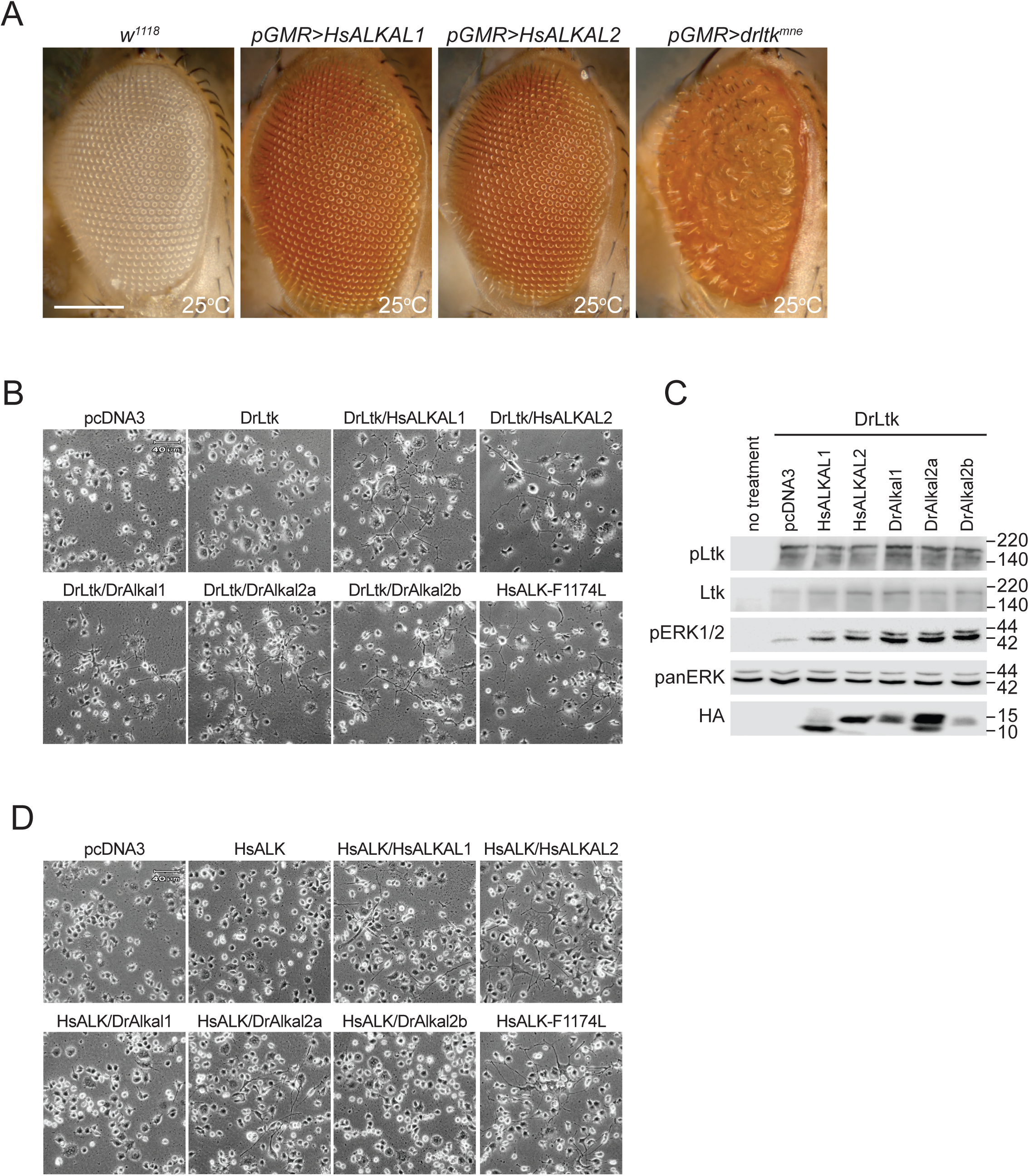
Activation of HsALK and DrLtk by the ALKAL proteins. (A) Wildtype fly eyes (*w^1118^*) show an organized ommatidia structure. Similarly, expression of either HsALKAL does not lead to a rough eye phenotype. The gain-of-function mutant DrLtk^mne^ was employed as positive control. (B) DrLtk is activated by both human and zebrafish ALKAL proteins, as shown by neurite outgrowth in PC12 cells transfected with both the ligand and the receptor as indicated. Gain-of-function HsALK-F1174L was employed as positive control. (C) Immunoblot for HEK293 cells treated with conditioned media. (D) Expression of HsALK in PC12 cells drive neurite outgrowth when coexpressed with human ALKALs and DrAlkal2a, but not DrAlkal1 or −2b. HsALK-F1174L was employed as positive control.

